# Ion mobility mass spectrometry unveils global protein conformations in response to conditions that promote and reverse liquid-liquid phase separation

**DOI:** 10.1101/2022.12.21.521395

**Authors:** Christina Glen Robb, Thuy P. Dao, Jakub Ujma, Carlos A. Castañeda, Rebecca Beveridge

## Abstract

Liquid-liquid phase separation (LLPS) is a process by which biomacromolecules, particularly proteins, condense into a dense phase that resembles liquid droplets. Dysregulation of LLPS is implicated in disease, yet the relationship between protein conformational changes and LLPS remain difficult to discern. This is due to the high flexibility and disordered nature of many proteins that phase separate under physiological conditions, and their tendency to oligomerise. Here we demonstrate that ion mobility mass spectrometry (IM-MS) overcomes these limitations. We used IM-MS to investigate the conformational states of full-length ubiquilin-2 (UBQLN2) protein, LLPS of which is driven by high salt concentration and reversed by noncovalent interactions with ubiquitin (Ub). IM-MS revealed that UBQLN2 exists as a mixture of monomers and dimers, and that increasing salt concentration causes the UBQLN2 dimers to undergo a subtle shift towards extended conformations. UBQLN2 binds to Ub in 2:1 and 2:2 UBQLN2:Ub complexes which have compact geometries compared to free UBQLN2 dimers. Together, these results suggest that extended conformations of UBQLN2 are correlated with UBQLN2’s ability to phase separate. Overall, delineating protein conformations that are implicit in LLPS will greatly increase understanding of the phase separation process, both in normal cell physiology and disease states.

## Introduction

Intrinsically disordered proteins (IDPs) exist and function without the fixed tertiary structure that was once thought to be required for all proteins to carry out their physiological roles. Instead, IDPs populate a wide range of conformations, from compact to extended, and rapidly interconvert between these various geometries, largely unhindered by energetic constraints [1]. IDPs and intrinsically-disordered regions (IDRs) in proteins are an important focus of research due to their high abundance, with 30% of the human proteome predicted to be disordered [2]. Dysregulation of the biophysical properties of IDPs is often associated with diseases such as cancer and neurodegenerative disorders, as IDPs are significantly involved in cell signalling networks [3, 4]. Additionally, long IDRs are often involved in multivalent interactions that contribute to liquid-liquid phase separation (LLPS), hypothesised to underlie formation of biomolecular condensates [5-7]. LLPS is important in normal cell physiology, and dysfunctional LLPS is involved in diseases such as amyotrophic lateral sclerosis (ALS) [8].

The ubiquitin (Ub)-binding proteasomal shuttle Ubiquilin-2 (UBQLN2) is an ALS-linked, IDR-containing protein that undergoes LLPS under physiological conditions [9-12]. LLPS of UBQLN2 is modulated by multivalent interactions among IDRs that include the STI1-II region and disease-associated PXX domain [11-14], as well as the folded N-terminal Ub-like (UBL) and C-terminal Ub-associating (UBA) domains that bind proteasomal subunits [15] and Ub/polyUb chains [16], respectively (Figure 1a, b). The STI1-II region drives UBQLN2 oligomerisation that is a prerequisite for its phase separation [10, 12], which occurs in response to increases in salt concentration and temperature [17]. Importantly, noncovalent interactions between monoUb and the UBA domain of UBQLN2 inhibits LLPS droplet formation by disrupting the protein-protein interactions that drive the process [10]. We hypothesise that the promotion or inhibition of UBQLN2 LLPS, which is sensitive to salt concentration, temperature, and Ub-binding, result from changes in the global UBQLN2 conformation. However, UBQLN2 conformations under these different conditions remain elusive to most biophysical techniques (small angle X-ray scattering (SAXS), analytical ultracentrifugation (AUC), and nuclear magnetic resonance (NMR)) as these methods require high protein concentrations at which UBQLN2 phase separates. Furthermore, the combination of high IDR content and oligomerisation propensity of UBQLN2 complicates interpretation of experiments that inform on conformation [10, 11].

**Figure 1.**
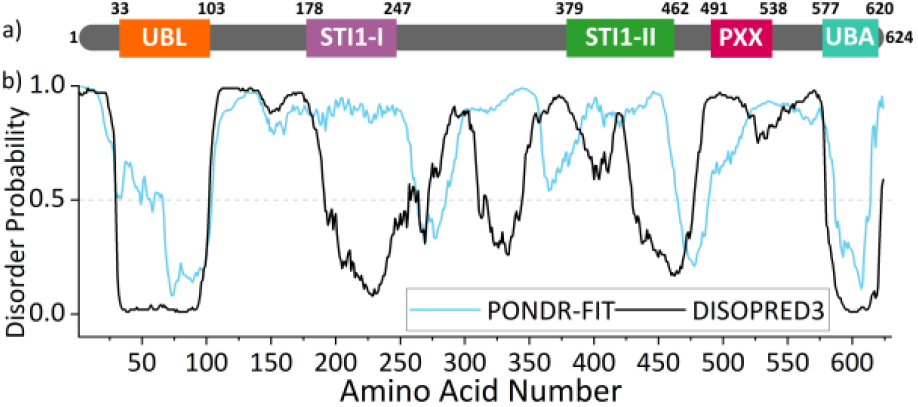
(a) Domain architecture of UBQLN2. (b) PONDR-FIT and DISOPRED3 predictions of disorder for UBQLN2, showing that the UBL and UBA are structured, with the region between being largely disordered. Figure adapted with permission from Dao et al. [10] Copyright 2018 Elsevier

Native mass spectrometry (nMS) and ion mobility mass spectrometry (IM-MS) have emerged as versatile and informative methods to study IDPs and proteins containing IDRs [18, 19]. Upon “soft” transfer from solution into the gas phase by nanoelectrospray ionisation (nESI), non-covalent interactions within and between proteins can be preserved, enabling the mass-to-charge ratio (*m/z*) measurement of intact proteins and protein complexes. Deconvolution of nMS data reveals the stoichiometric composition of a protein complex (*via* mass, *m*), and the degree of compaction or extension of a protein/protein complex can be correlated with a number of associated charges, *z* [20]. Compact conformations have a limited solvent accessible surface area on which protons can be accommodated, and hence have a low number of charges, whereas extended conformations have space for a large number of protons and therefore have a high charge state [21]. The wide charge state distribution, which is a hallmark of IDPs in nMS experiments, reflects their wide range of adopted conformations. Proteins are generally ionised from ammonium acetate (AmAc) solution, which is volatile and evaporates readily from the protein during desolvation. Many salts and buffers cause significant adduction to the protein ions, causing challenges in spectral assignment by broadening the peaks and lowering the signal intensity. Several approaches have been developed towards allowing proteins to be ionised from commonly used buffers such as phosphate buffered saline (PBS), including the use of small nanoelectrospray emitters [22] or adding ammonium acetate to the buffer of choice [23]. Towards analysing proteins from more physiological conditions, Sharon and co-workers have shown considerable success with nMS of overexpressed proteins in crude cell lysates [24]. In addition to nMS, hydrogen-deuterium exchange-MS can also reveal the effect of salts, metal ions and buffers on the metastable conformations of IDPs [25, 26].

The hybrid method of IM-MS adds an extra dimension of analysis by separating protein ions on the basis of their overall size, meaning that multiple conformations can be separated from one *m/z* ratio. Larger conformations travel slower through an IM drift region, due to an increased number of collisions with gas molecules [27]. The experiment yields arrival time distributions (ATDs) which can be converted to rotationally averaged collisional cross sections (CCSs) to report on the size of the proteins of interest in units of nm^2^ [28]. Whilst CCSs can be directly calculated from ATDs with the use of drift tube IM-MS apparatus, the use of travelling wave IM-MS, as used in this study, requires measuring appropriate calibrant proteins with known CCS values. IM-MS therefore provides information on the range of sizes that a protein can exist in, which infers on their conformational heterogeneity and hence their dynamic behaviour [29]. Advantages of using IM-MS to study IDPs include its ability to measure the size range of every species that is present in a stoichiometric mixture, including co-existing conformations of the same species, without any ensemble averaging and without interference from other complexes that are present in the mixture. Low protein concentrations are required, and the method is equally suited to compact and extended conformations and does not favour the folded state of a protein [20]. nMS and IM-MS have been widely employed to study disordered protein systems such as p27 [21], α-synuclein [30] and melanoma-associated antigen A4 (MAGE-A4) [31]. IM-MS was also recently used to study the conformations of phase-separating proteins Fused in Sarcoma (FUS) and Transactive Response DNA-binding protein (TDP-43) in response to differing pH of the solution from which they were analysed [32].

In this work, we used IM-MS to measure the conformational distributions of UBQLN2 in its soluble form, delineate its conformational response to increased salt concentration that drives LLPS, and interrogate the complexes it forms with Ub which reverses and inhibits LLPS. We identified that UBQLN2 exists as a mixture of monomers and dimers, both with an extraordinarily wide range of conformations. At increased salt concentration, the dimers undergo a subtle shift to more extended conformations which we hypothesise are implicit in driving LLPS. In contrast, the presence of Ub stabilises compact conformations of UBQLN2 dimers which we propose are unable to form the multivalent intermolecular interactions required for LLPS. These findings were enabled by the ability of IM-MS to (i) measure full-length UBQLN2 proteins, (ii) report on the conformations of individual complexes that are present in a mixture, and (iii) perform measurements at protein concentrations below the threshold for LLPS. These IM-MS experiments therefore reveal conformational changes associated with LLPS whilst retaining UBQLN2 in its soluble state.

## Methods

### Materials

Ammonium acetate solution (AmAc) was prepared at pH 6.8 from ultra-pure water (18.2 MΩ.cm, Millipore) and analytical grade ammonium acetate solid. (Fisher Scientific).

### Expression and purification of UBQLN2 and Ub

E. coli NiCo21 (DE3) (New England BioLabs) cells containing full length (FL)-UBQLN2, UBQLN2-ΔUBA and ubiquitin (Ub) in pET24b (+) plasmid (Novagen) were grown in Luria-Bertani (LB) broth at 37°C to an optical density at 600 nm of 0.6-0.8. Expression was then induced with β-D-1-thiogalactopyranoside (IPTG) to a final concentration of 0.5 mM at 37 °C overnight. Bacteria were pelleted, frozen, and lysed in pH 8 buffer containing 50 mM Tris, 1 mM EDTA, 1 mM PMSF, 4 mM MgCl_2_, 0.5 mg/mL lysozyme and Pierce Universal Nuclease (ThermoFisher). For full-length UBQLN2 and UBQLN2-ΔUBA, NaCl was added to the cleared lysate at 30° C to the final concentrations of 0.5 M and 1 M, respectively, to induce UBQLN2 phase separation. UBQLN2 droplets were pelleted by centrifugation and then resuspended in pH 6.8 buffer containing 20 mM NaPhosphate, 0.5 mM EDTA, 0.1 mM TCEP, 0.02% NaN_3_. Leftover NaCl was removed by passing the solution through HiTrap desalting column (GE Healthcare). The UBQLN2 purification protocol here is adapted from [33]. For Ub, perchloric acid was added to the lysate to a final pH of about 1. Precipitated proteins were cleared by centrifugation. The supernatant was diluted 1:1 with 50 mM AmAc, pH 4.5, loaded onto HiTrap SP HP column (GE Healthcare), and eluted with a gradient to 1 M NaCl in 50 mM AmAc, pH 4.5. Fractions containing Ub were concentrated and buffer exchanged into pH 6.8 buffer (see above) using Vivaspin 6 centrifugal concentrators with a molecular weight cutoff (MWCO) of 5000 Da (Sartorius). Purified proteins were frozen at −80°C.

### Preparation of proteins for nMS and IM-MS

Full length (FL)-UBQLN2 (30 μM) and UBQLN2-ΔUBA (45 μM) were buffer exchanged into 10 mM AmAc pH 6.8 using 96-well Microdialysis plates (Thermo Fisher Scientific, Waltham, MA USA). Ub was buffer exchanged using Bio-Rad Micro Bio-Spin P6 columns (Bio-Rad, Hercules, CA, USA). Final protein concentrations were determined using a NanoDrop spectrophotometer (Thermo Fisher Scientific Waltham, MA USA) using the A280 method. Protein concentrations were calculated with extinction coefficients of 11460 and 1490 M^-1^ cm^-1^ for the UBQLN2 constructs and Ub, respectively.

### Native mass spectrometry and ion mobility

FL-UBQLN2 was diluted to a protein concentration of 15 μM and final AmAc concentrations of 10 mM, 50 mM, or 75 mM (pH 6.8) for experiments on the unbound protein. UBQLN2-ΔUBA was diluted to 5 μM, and final AmAc concentrations of 10 mM, 55 mM and 100 mM (pH 6.8). For experiments with FL-UBQLN2 and Ub, a 1:4 molar ratio of FL-UBQLN2 monomer to Ub was mixed by adding 15 μM FL-UBQLN2 (10 mM AmAc) to an equal volume of 60 μM Ub (10 mM AmAc). All samples were allowed to equilibrate on ice for at least 30 minutes.

Ion mobility mass spectrometry data was acquired on a Waters Synapt G2-Si (Waters Corporation, Wilmslow, UK) instrument with an 8k quadrupole operated in “Sensitivity” mode. Proteins were subject to nanoelectrospray ionisation (nESI) in positive mode with a nanospray tip pulled in-house with a Flaming/Brown P-97 micropipette puller from thin-walled glass capillaries (i.d. 0.78 mm, o.d. 1.0 mm, 10 cm length, both from Sutter Instrument Co., Novato, CA, USA). A positive potential of 1.2-1.6 kV was applied to the solution via a thin platinum wire. Other non-default instrument settings include: sampling cone voltage 60V, collision voltage 5V, trap gas flow 3.5-4 ml/min, source temperature 40 °C. Ion mobility data of FL-UBQLN2 alone and UBQLN2-ΔUBA were collected at travelling-wave velocity of 400 m/s and height of 40V. Ion mobility data of UBQLN2:Ub complexes were collected at travelling-wave velocity of 325 m/s and ramped height of 25-40V. Helium and nitrogen (IMS) gas flows were 180 ml/min and 90 ml/min for experiments involving FL-UBQLN2 and UBQLN2-ΔUBA, and 150 and 75 ml/min for experiments involving FL-UBQLN2:Ub complexes. Instrument was allowed to settle for one hour prior to experiments. A manual quadrupole RF profile was applied to improve transmission of ions from *m/z* 2750 and upwards. CCS calibration was performed using IMSCal19 (Waters Corporation, Wilmslow, UK) [28, 34] with β-lactoglobulin and bovine serum albumin used as CCS calibrants [35].

### Data processing

Mass spectra were initially processed in MassLynx v4.2 (Waters Corporation, Wilmslow, UK). IM-MS profiles were created in OriginPro 2022 (OriginLab Corporation, Northampton, MA, USA) by extracting ion mobility spectra from selected charge states in the mass spectrum, then normalising and averaging spectra collected from multiple tips, on the same day. For FL-UBQLN2, normalised data from across multiple days was then averaged and the standard deviation across multiple days was calculated using Descriptive Statistics function in OriginPro 2022 and reported as error bars. For UBQLN2-ΔUBA, data was collected in triplicate on one day and the standard deviation across measurements was calculated using Descriptive Statistics function in OriginPro 2022 and reported as error bars. Peak fitting was performed on the average profiles using Multiple Peak Fit function: Gauss peak fitting in OriginPro 2022. Peak apexes and widths were conserved between salt concentrations to enable tracking of relative abundances, and iterations were performed until an R2 > 0.98 was achieved whilst simultaneously avoiding overfitting of peaks. 3 peaks were fitted to FL-UBQLN2, whilst the data for UBQLN2 ΔUBA appears to be composed of 4 components/species, thus 4 peaks were fitted. % area of each peak was calculated by dividing the area of a peak by the sum of all areas. Red dashed line represents the cumulative fit for each profile.

For Figure S11, baseline subtraction was performed using Peak Analyser: Create baseline function in OriginPro2022 where anchor points along baseline were selected manually and connected using BSpline interpolation method. Text files generated from CCS calibration using IMSCal19 were collated in Microsoft Excel and scatter plots of CCS versus charge were plotted in OriginPro2022 by extracting the maxima(s) for each charge states’ CCS values. Figures were created using Inkscape 1.2.1 (inkscape.org).

### Bright-field imaging of phase separation

Samples were prepared to contain 15 μM FL-UBQLN2 or 5-10 μM UBQLN2-ΔUBA and different concentrations of AmAc, pH 6.8 from 10 mM to 300 mM (FL-UBQLN2) and 10 to 400 mM (UBQLN2-ΔUBA) and incubated on ice. Samples were added to Eisco Labs Microscope Slides, with Single Concavity, and covered with MatTek coverslips that had been coated with 5% bovine serum albumin (BSA) to minimise changes due to surface interactions, and incubated coverslip-side down at 20 ºC for 10 min. Phase separation was imaged on an ONI Nanoimager (Oxford Nanoimaging Ltd, Oxford, UK) equipped with a Hamamatsu sCMOS ORCA flash 4.0 V3 camera using an Olympus 100×/1.4 N.A. objective. Images were prepared using Fiji [36] and FigureJ plugin.

## Results and discussion

### Delineating the effect of salt concentration on the mass spectrometry profile of FL-UBQLN2

As increased salt concentration promotes UBQLN2 LLPS, our first objective was to determine how changing the salt concentration affects the conformational ensemble of FL-UBQLN2 by spraying the protein from ammonium acetate (AmAc) concentrations of 10 mM, 50 mM and 75 mM representing low, medium, and high salt concentrations, respectively. Whilst full length (FL)-UBQLN2 does not phase separate under these conditions, AmAc is capable of inducing LLPS at higher concentrations (Figure S1). When analysed from 10 mM AmAc (Figure 2a), FL-UBQLN2 exists as a mixture of monomers and dimers with the monomer present in charge states 16+ to 75+ (Δ*z*=59) and the dimer present in charge states 22+ to 83+ (Δ*z*=61). The measured mass of the monomer is 65 640 Da and that of the dimer is 131 280 Da. Both charge state distributions represent extremely dynamic species, as the maximum Δz of a structured protein was previously found to be 6 for proteins up to 150 kDa in mass [20]. We focus primarily on the dimeric species, as these undergo changes in response to solution conditions and we therefore assign these as being implicit in LLPS. We assign the low-charged dimers of charge states 22+ to 31+ (*m/z* 4200-5750) as compact, dimer charge states 32+ to 43+ (*m/z* 3000-4200) as being intermediate, and dimer charge states 44+ and above (*m/z* 750-3000) as being extended. We assign the cause of the raised baseline to the width of the *m/z* peaks due to isotope distributions, incomplete desolvation and modifications such as methionine oxidation that broaden the width of each protein peak. At medium and high AmAc concentrations (Figure 2b and c) the compact dimers have a much lower signal intensity. This suggests that increased salt concentration, which drives UBQLN2 LLPS, causes a decrease in the abundance of compact dimers. Of note, 75mM was the highest concentration of AmAc from which the protein could be analysed without the nanoelectrospray needle becoming blocked by insoluble protein. Fully annotated versions of the spectra shown in Figure 2 can be seen in Figure S2.

**Figure 2.**
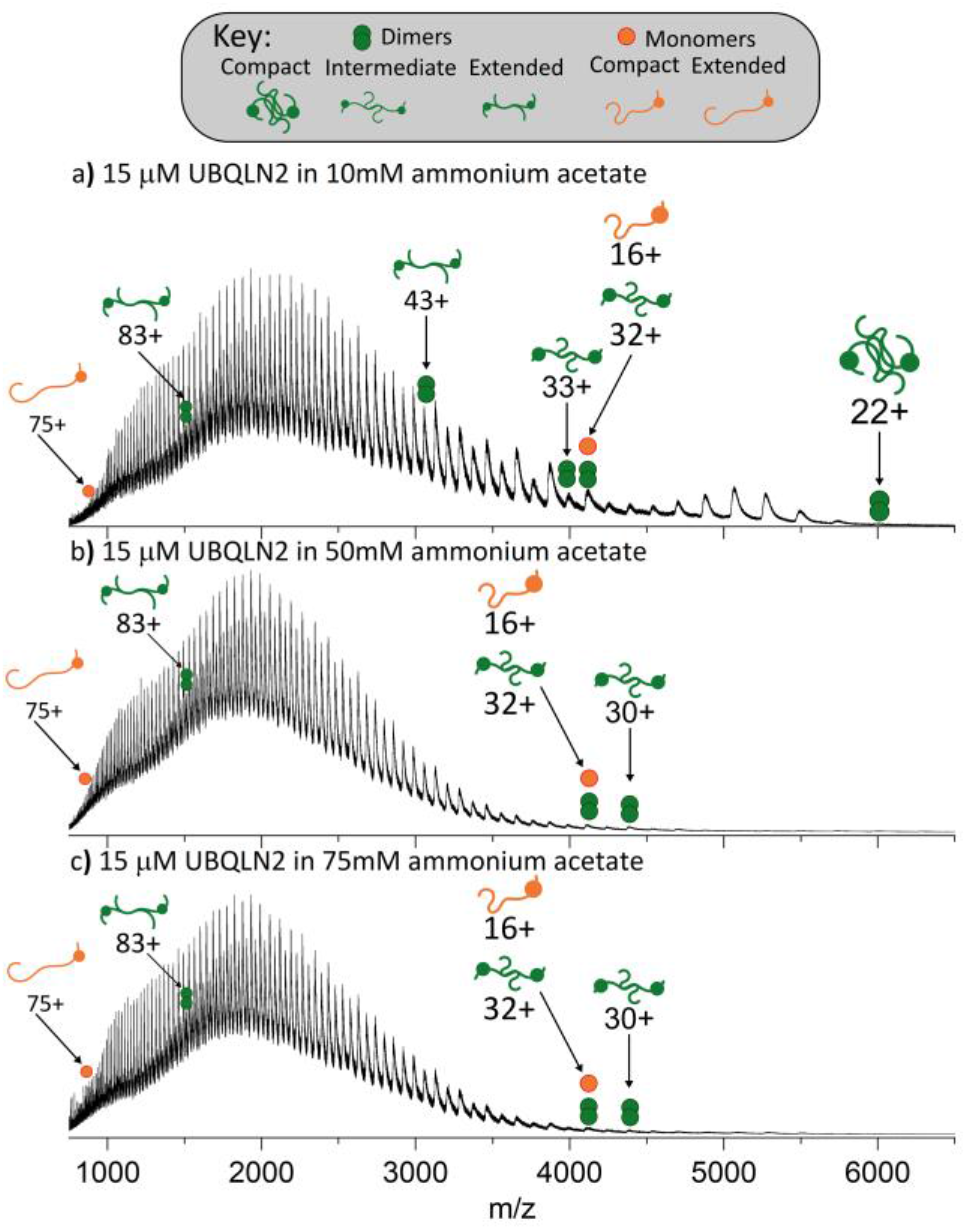
nMS reveals a loss of compact UBQLN2 dimers with increasing salt concentration. FL-UBQLN2 (15 μM monomer) is analysed from AmAc concentrations of 10 mM (a), 50 mM (b) and 75 mM (c). Fully annotated spectra can be seen in Figure S2, and *m/z* ratios for the UBQLN2 monomer and dimers are given in Tables S1 and S2, respectively.

A construct of UBQLN2 lacking the UBA domain (UBQLN2-ΔUBA), which undergoes LLPS to a much lower degree than FL-UBQLN2 [10], was also subject to nMS analysis. After ascertaining that UBQLN2-ΔUBA remains soluble in AmAc concentrations of up to 400 mM (Figure S3), we analysed UBQLN2-ΔUBA from AmAc concentrations of 10 mM, 55 mM and 100 mM (Figure S4). This construct follows the same trend as FL-UBQLN2 in that low charge states of the dimer (25+ to 36+), corresponding to compact dimeric conformations, are represented to a much lower extent at higher AmAc concentrations (Figure S4). Therefore, these UBQLN2-ΔUBA data are consistent with the elongation of FL-UBQLN2 observed with increased salt concentration. This strengthens the hypothesis that UBQLN2 elongation is an intermediate step towards LLPS, rather than what remains soluble after LLPS of a different conformation has taken place.

### Ion mobility mass spectrometry reveals elongation of intermediate-charged dimer conformations at increasing salt concentrations

IM-MS was subsequently used to investigate whether further conformational changes of UBQLN2 can be detected upon increasing the concentration of AmAc in the starting solution. The high charge states corresponding to elongated conformations remain unaffected (47+ dimer, Figure S5). While the compact conformations couldn’t be compared as the signal intensity for medium and high salt concentrations were too low, arrival time distributions (ATDs) shown for charge states 24+ to 30+ in 10mM AmAc (Figure S6) confirm that FL-UBQLN2 at charge states 24+ to 27+ are mainly compact, and then become more elongated between 28+ and 30+. The intermediate-charged dimer populations underwent a subtle shift towards more elongated conformations at high salt concentration, represented by the peak corresponding to the 33+ dimer (*m/z* 3975, Figure 3a) of FL-UBQLN2, as described below.

**Figure 3.**
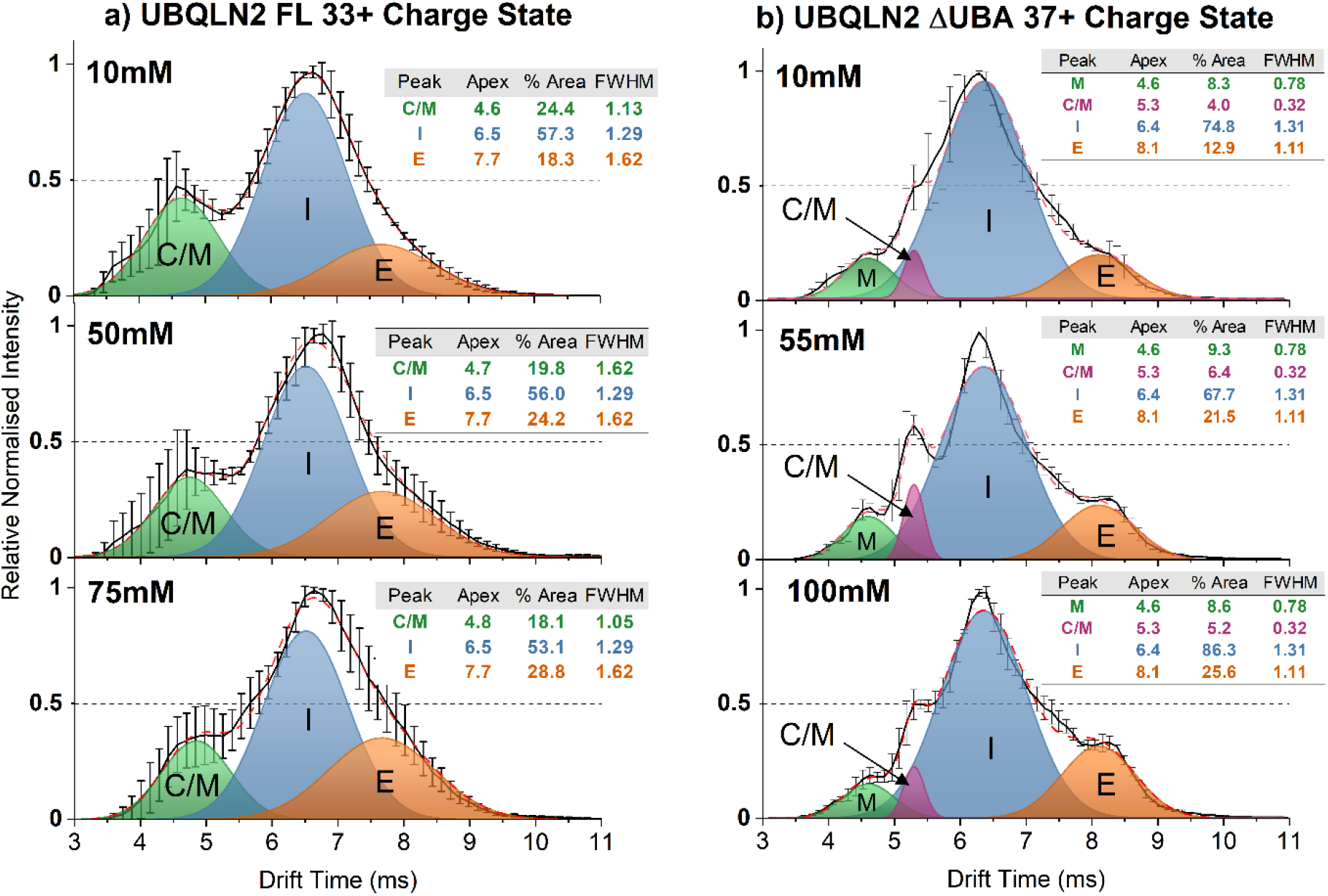
IM-MS shows that the intermediate-charged dimer conformations of FL-UBQLN2 and UBQLN2-ΔUBA become larger when analysed from high salt concentrations. (a) IM spectra of the FL-UBQLN2 33+ dimer analysed from AmAc concentrations of 10 mM (top), 50 mM (middle) and 75 mM (bottom). Gaussian curves represent populations corresponding to intermediate dimers (I), extended dimers (E), and overlapping monomer/compact dimer (C/M). Solid black lines in figure 3a are the average of 2-3 measurements across three days (n = 7 − 9) and error bars represent standard deviation across days. (b) IM spectra of the UBQLN2-ΔUBA 37+ dimer analysed from AmAc concentrations of 10 mM (top), 55 mM (middle) and 100 mM (bottom). Gaussian curves correspond to monomers (M), overlapping monomers and compact dimers (C/M), intermediate dimers (I) and extended dimers (E). Solid black line is the average of n=3 measurements taken on one day and error bars represent standard deviation across measurements. Dotted red line is the sum of the Gaussian fits.

The ATDs of the FL-UBQLN2 33+ dimer sprayed from low, medium, and high AmAc concentrations are shown in Figure 3a, and Gaussian curves were fitted to the data to represent conformational populations. Curves labelled I and E correspond to intermediate and extended conformations of the 33+ dimer, respectively, and the relative areas of the extended conformation are 18.3% (10 mM AmAc), 24.2% (50 mM AmAc), and 28.8% (75 mM AmAc). Overall, this shows that there is a higher abundance of more elongated conformations at 75 mM AmAc than at lower concentrations, which is the condition closest to that at which LLPS occurs. The peak denoted C/M corresponds to an overlapping population of monomer and compact dimer that have coincident arrival times and m/z ratios due to ‘tailing’ of the adjacent 16+ monomer (Figure S7). The relative areas of the C/M curves are 24.4% (10 mM AmAc), 19.8% (50 mM AmAc) and 18.1% (75 mM AmAc). While it is uncertain what proportion of the signal in this region arises from either species, ATDs for the overlapping 32+ dimer/16+ monomer are shown in Figure S8 which shows that the conformation of the 16+ monomer is largely unaffected by increasing AmAc concentrations. The absence of oligomers of higher order than a dimer is shown in the m/z vs. drift time plot in Figure S7, in which profiles corresponding only to monomers and dimers can be observed.

ATDs of the 37+ charge state of UBQLN2-ΔUBA were also compared when sprayed from 10mM, 55mM and 100mM AmAc (Figure 3b). Species with the earliest arrival times are again attributed to ‘tailing’ of the monomer peaks (Figure S9) and are represented by Gaussian curves labelled monomer (M) and overlapping monomers and compact dimers (C/M), while the dimers are represented by curves corresponding to the intermediate dimer (I) and the extended dimer (E). Here, the relative area of the extended conformation increases from 12.9% at 10 mM AmAc to 21.5% at 55 mM AmAc and 25.6% at 100 mM AmAc. Again, the extended conformation increases in intensity as AmAc concentration increases and is also becoming more distinct from the intermediate conformation. This is best observed in Figure S10, which shows the ATDs without the Gaussian curves. We envision that an extended conformation of FL-UBQLN2 is also becoming more intense at high AmAc, but in this case the conformation is not resolvable in the ATD due to increased dynamics of the system.

### UBQLN2 binds to Ub in 2:1 and 2:2 complexes which stabilises compact conformations

As specific interactions between UBQLN2 and Ub (monoUb) drive disassembly of UBQLN2 biomolecular condensates [10], we used nMS to investigate the stoichiometry and conformation of the FL-UBQLN2:Ub complexes. The native mass spectrum of a 1:4 molar ratio of UBQLN2 monomer to Ub reveals that Ub binds to UBQLN2 dimers with either 2:1 or 2:2 UBQLN2 to Ub stoichiometry (Figure 4). The measured mass of the 2:1 complex is 139 820 Da and that of the 2:2 complex is 148 440 Da. Charge states for the 2:1 complex, labelled with green dotted lines, range from 22+ to 47+, with charge states 30+ to 35+ being much lower in intensity and not resolved from other peaks in the same *m/z* range (Figure S10). Complexes with 2:2 stoichiometry labelled with orange dotted lines, range from 24+ to 49+ with a gap in resolvable peaks from 33+ to 39+, as well at 41+ and 43+, and are lower in intensity than the 2:1 complexes. Whilst monomeric UBQLN2 is not observed to bind to Ub, this cannot be ruled out due to the overlapping *m*/*z* values between the 1:1 complex and evenly-charged 2:2 complex. However, no increase in abundance of the evenly charged 2:2 complexes is observed compared to the oddly charged, which would be the indication of 1:1 complex. Moreover, signal corresponding to monomeric UBQLN2 remains at high intensity in the mixture with Ub, indicating that it is still present in solution, whereas the signal intensity for unbound UBQLN2 dimers is depleted as it complexes with Ub.

**Figure 4:**
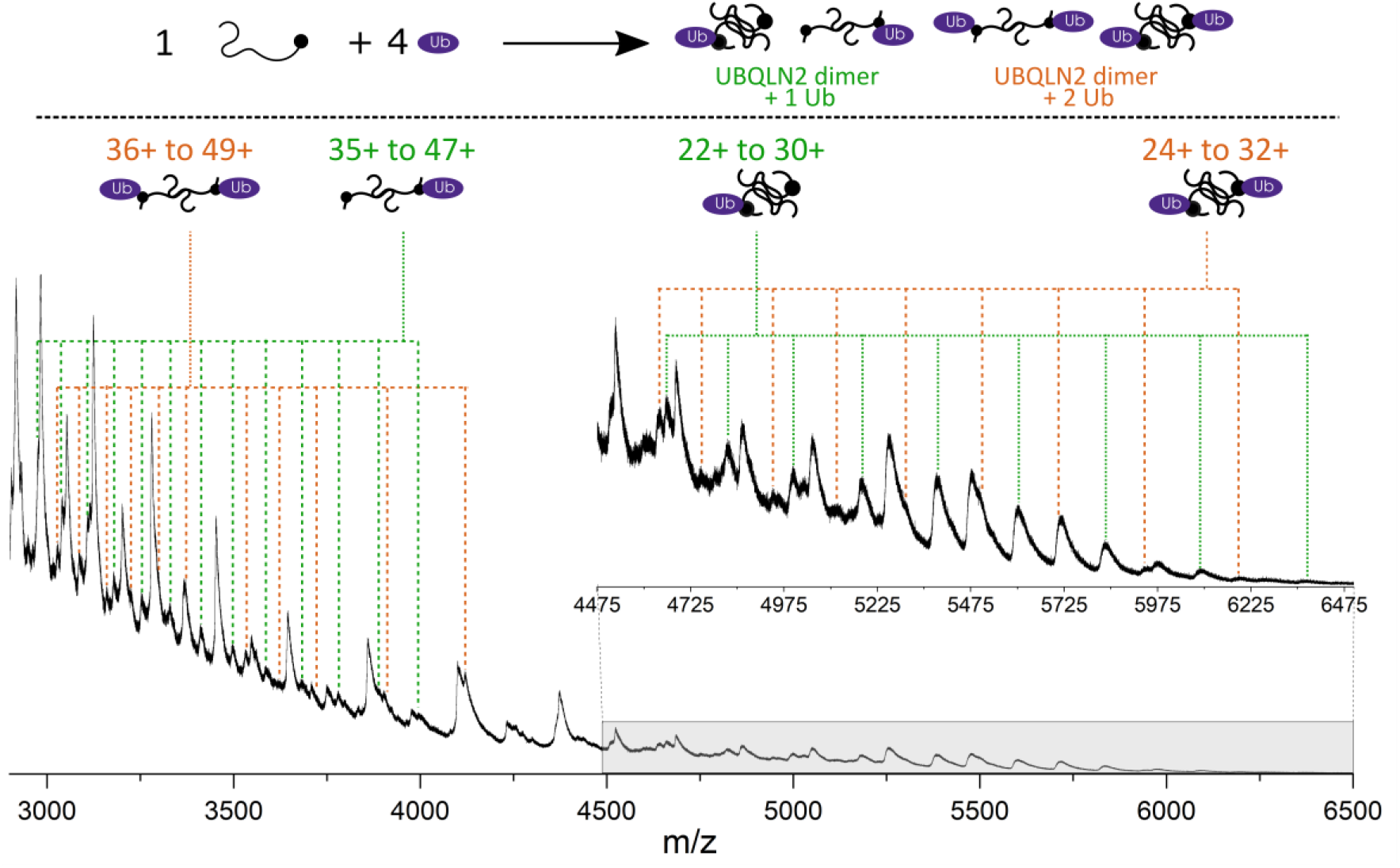
Native mass spectrum (2900-6500 *m/z*) resulting from a 1:4 molar ratio of FL-UBQLN2:Ub in 10 mM AmAc. Mixture results in the formation of 2:1 (green) and 2:2 (orange) FL-UBQLN2 to Ub complexes. *m/z* units for the 2:1 and 2:2 UBQLN2:Ub complexes are given in Tables S3 and S4, respectively.

The charge state range of both Ub-containing complexes (Δ*z* = 25 in both cases) is much narrower than the range observed for FL-UBQLN2 alone (Δ*z* = 61) but is still very wide compared to a structured protein of a similar size (Δ*z* = 6 for Serum Amyloid P Pentamer, 128 kDa [20]), suggesting the complexes are still highly dynamic. A comparison of the nMS of UBQLN2 and UBQLN2:Ub complexes can be seen in Figure S11. Another feature of the mass spectrum worth noting is the reduction in intensity of the peaks previously assigned as intermediate-charged UBQLN2 dimers (3000-4200 *m/z*), indicating that these are the dimeric charge states that are binding to Ub and are therefore of lower abundance in solution. These also correspond to the charge states which form elongated conformations in response to salt concentration (Figure 3). We hypothesise these are the conformations that are responsive to variations in solution conditions and mediate LLPS of UBQLN2.

As mentioned previously, it is possible to convert drift time values collected in IM-MS experiments to rotationally averaged collisional cross sections (CCS) using calibrant proteins with known CCS values [28, 34]. This has been performed for UBQLN2 and UBQLN2:Ub complexes to allow comparison of sizes between different species.

Figure 5a shows the CCS values for individual charge states of UBQLN2 dimers and 2:1 and 2:2 UBQLN2:Ub complexes. For the UBQLN2 dimers, only odd charge states are shown above 31+ to avoid interference of signal from the monomer. For UBQLN2:Ub complexes, charge states which are not calibrated are marked with red crosses in figure S11. Charge states 22+ to 25+ exist in a single compact conformational family around 68 nm^2^, while charge states 26+ to 41+ exist in two conformational families shown by a bimodal ATD for these charge states (Figure 5b). Charge states 43+ to 85+ correspond to a conformational family that increases in size with increasing charge, typical of IDPs. Overall, UBQLN2 dimers range from 68 to 260 nm^2^, which is a huge range of conformations for a complex of this mass. For reference, the Serum Amyloid P Pentamer (128 kDa) has a CCS range of 59-64 nm^2^ [20].

**Figure 5:**
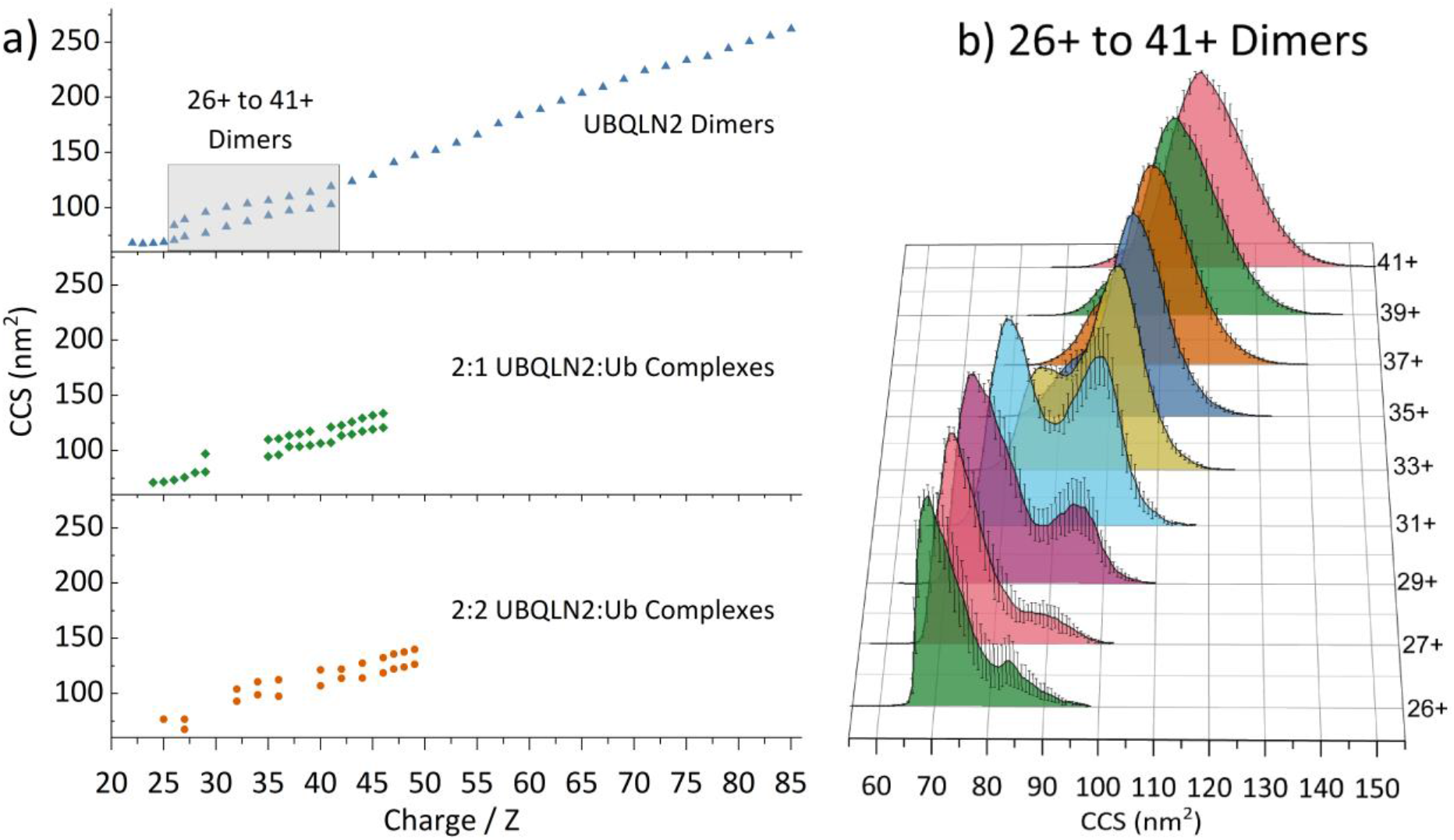
a) Collision cross section distributions of UBQLN2 dimers, UBQLN2:Ub 2:1 complexes and 2:2 complexes. b) Calibrated collisional cross section distributions for free UBQLN2 dimer charge states 26+ to 41+. A bimodal distribution can be observed wherein the compact conformation dominates from 26+ to 31+ until a conformational switch occurs and the more elongated conformation dominates from 35+, trending larger as charge increases. Solid black line represents average from three measurements and error bars represent standard deviation across three measurements.

UBQLN2:Ub complexes are present in a smaller range of CCSs, indicating that Ub stabilises compact conformations of UBQLN2, with the largest conformation in both cases being ∼135 nm^2^ as opposed to 260 nm^2^ for the UBQLN2 dimer. For the 2:1 UBQLN2:Ub complex, a compact conformational family exists from 71-80 nm^2^ (charge states 24+ to 29+). 29+ has two conformational families at 80 and 98 nm^2^, then there is a gap in resolvable *m*/*z* peaks from 30+ to 34+ (Figure S11, red crosses). From 35+ to 47+ the large conformation increases in size gradually with CCSs from 110-132 nm^2^, whilst the smaller conformation of these charge states has slightly larger changes in CCS between charge states 36+ to 37+ and 42+ to 43+.

The 2:2 UBQLN2:Ub complexes follow a similar trend, in which compact complexes exist at CCSs of 68-76 nm^2^ (*z* = 25+ to 27+), there is then a gap in resolvable *m/z* peaks (CCS 78-92 nm^2^, *z* = 27+ to 31+) and from *z* = 32+ to 49+ there are two conformations per charge state, with an overall increase in CCS with respect to charge. Not shown in this region are charge states *z* = 37+ to 39+, CCS 107-112 nm^2^, as they are poorly resolved and in low abundance. The largest CCS for the 2:2 complex is marginally larger than the 2:1 complex: 140 vs 135 nm^2^. We were interested to note that the most compact conformations of the UBQLN2:Ub complexes have similar CCS values to the most compact UBQLN2 dimers despite the increase in mass, suggesting that addition of ubiquitin is stabilising even more compact conformations of UBQLN2 than the free UBQLN2 dimers. Additionally, the complexes with Ub do not reach the size of the most extended conformations for free UBQLN2 dimers. When considering LLPS, this could mean addition of ubiquitin is preventing UBQLN2 inter-molecular interactions by stabilising compact conformations that are likely to be more spherical in shape than the free UBQLN2 dimers.

## Conclusions

In this work, we demonstrated the strength of nMS and IM-MS in elucidating conformational details of UBQLN2 in conditions where LLPS is promoted (high salt concentration) and inhibited (presence of Ub) (Figure 6). We found that LLPS-promoting high salt concentration depletes compact conformations and causes a subtle shift of UBQLN2 towards more elongated conformations. Conversely, Ub, which inhibits and reverses LLPS of UBQLN2, binds to and favours more compact conformations of UBQLN2 dimers.

**Figure 6:**
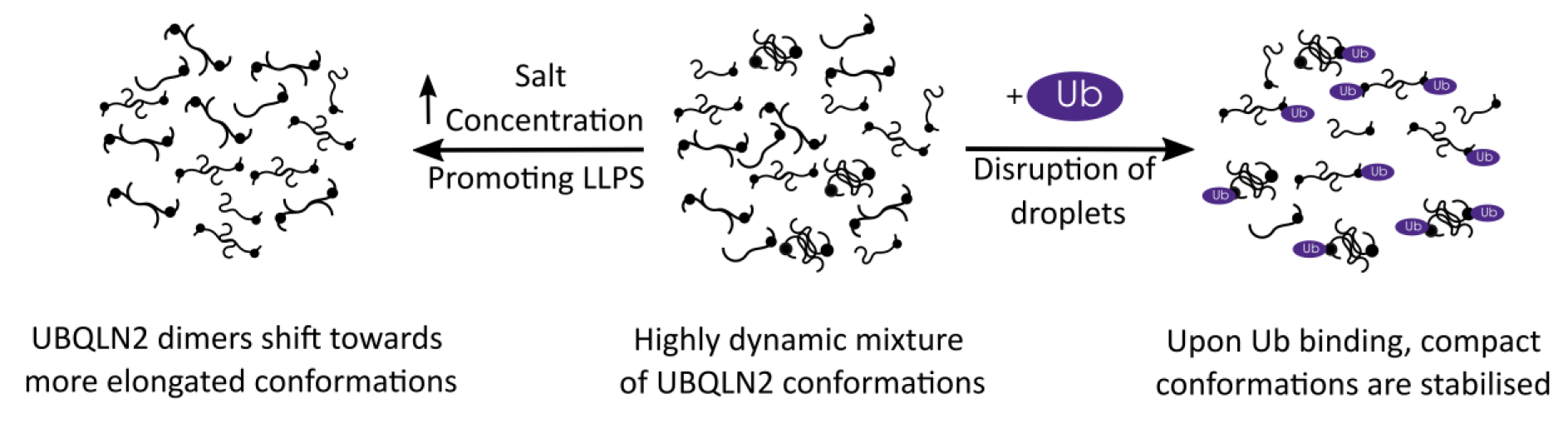
Model of changes in full-length UBQLN2 conformational state as a function of LLPS-promoting (salt addition) and LLPS-inhibiting (monoUb addition) conditions.

Determination of how conformation contributes to LLPS of specific proteins has been hampered by the challenges corresponding to alternative biophysical methods to measure conformational disparity. These include the size limitation in NMR experiments, the high protein concentrations required for methods such as NMR and AUC, and the inability of many methods to differentiate multiple complexes present in a mixture. We present nMS and IM-MS as versatile, label-free methods to measure individual complexes in a mixture. These methods reveal subtle shifts in protein conformations that result from alterations in the solution conditions from which they are analysed. A strong advantage of these MS methods is the sensitivity: proteins can be measured at low concentrations below the threshold for LLPS, thereby delineating conformational changes that occur *en route* to droplet formation.

The strength of MS in determining the underlying mechanisms of LLPS has been highlighted in two recent publications. Sahin et al. [32] used nMS and IM-MS to measure the conformations of engineered FUS and TDP-43 constructs that contain a spider silk domain for solubility. These proteins undergo LLPS at neutral pH but remain soluble at high pH, allowing the authors to track changes in conformation as they reduce the pH and move towards LLPS conditions. They observed that FUS undergoes an unfolded-to-globular transition as the pH is shifted from 12.5 to 7, which they attribute to conformational changes associated with LLPS, whereas TDP-43 oligomerises into partially disordered dimers and trimers. Ubbiali et al. [37] used crosslinking-MS to investigate the conformations of the IDP α-Synuclein under LLPS conditions, and discovered that α-Synuclein shifts towards more elongated conformations, making it amenable to interprotein interactions. This agrees with our assertions about UBQLN2, and we propose that elongation may be a common factor in the LLPS of IDPs, as it allows the formation of multivalent long-range interactions among protein molecules.

Our findings about the conformational status of UBQLN2 under several conditions have allowed us to speculate on how this affects the function of the protein *in vivo*. We propose that when UBQLN2 is in its free, unbound state, it is present in elongated conformations. This enables UBQLN2 to interact with other UBQLN2 molecules via the multivalent interactions involving the folded and disordered regions that promote its phase separation [10, 17]. Additionally, these interactions may include other protein components such as RNA-binding proteins that are found in stress granules and known to interact with UBQLN2 [38, 39]. Notably, this study investigates changes in UBQLN2 conformations in response to AmAc concentration, which we have shown induces LLPS in a similar manner to NaCl (Fig S1) [10]. Typical intracellular salts such as NaCl, KCl and MgCl_2_ may have subtly different effects on the conformation of IDPs [40-42], so it will be important to include these salts in future IM-MS studies towards bridging the gap with experiments in physiological conditions.

Upon binding ubiquitin, we speculate that interactions between UBQLN2 dimers are disrupted, resulting in more compact conformations of UBQLN2 that do not favour phase separation. These results are in line with prior studies that suggest UBQLN2 and other Ub-binding shuttle proteins undergo a change in conformation for their biological function [13, 43]. In this way, noncovalent interactions between UBQLN2 and monoubiquitinated substrates can potentially drive disassembly of UBQLN2 droplets or remove UBQLN2 from condensates inside cells. This research, in the future, will aid in determining how the conformational states of UBQLN2 are affected by ALS-linked mutations as well as engagement with other protein quality control components such as polyubiquitin chains and proteasomal receptors.

## Supplementary Information

Additional nMS, IM-MS and Brightfield microscopy data to support these results, as well as tables showing m/z units for all proteins and protein complexes analysed.

## Acknowledgements

RB is supported by a UKRI Future Leaders Fellowship (Grant Reference MR/T020970/1) and a Chancellor’s Fellowship from the University of Strathclyde. CGR acknowledges funding from Waters Corporation (UK). T.P.D. and C.A.C. are supported by NSF CAREER (MCB 1750462). We also acknowledge Magnus Kjaergaard and Alex Holehouse of IDPseminars for introducing the authors and initiating this fruitful collaboration!

---

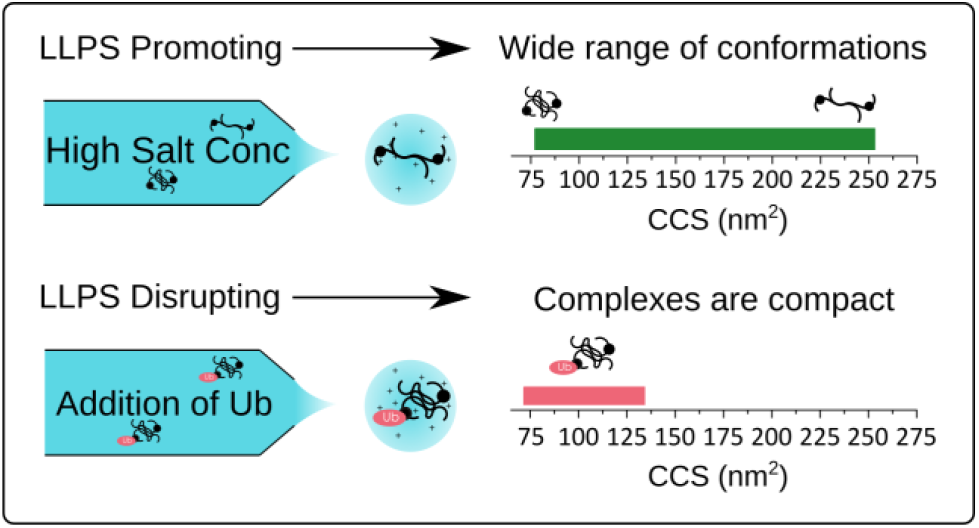

For table of contents only

