## Supplementary Information for "Ion mobility mass spectrometry unveils global protein conformations in response to conditions that promote and reverse liquid-liquid phase separation"

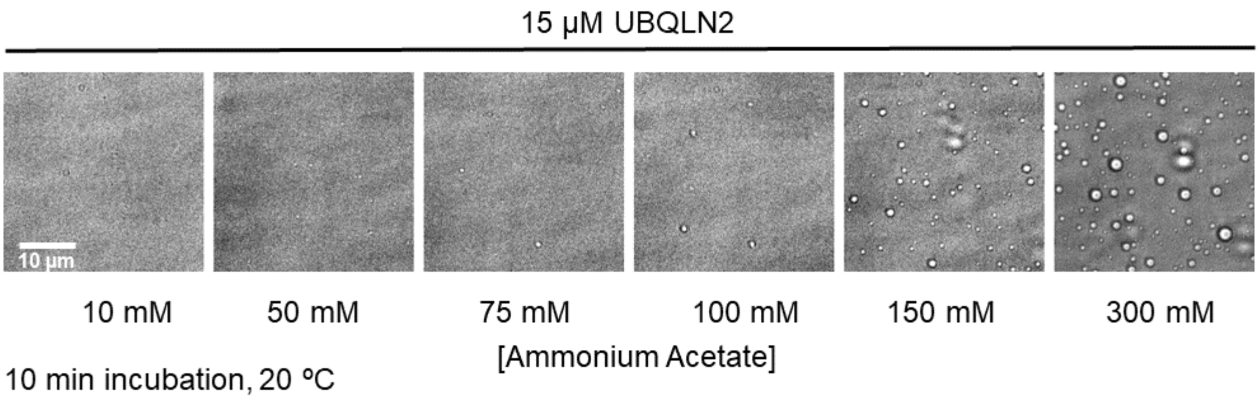


**Figure S1: Brightfield microscopy showing that FL-UBQLN2 is mostly soluble up to 100 mM ammonium acetate but undergoes LLPS at higher ammonium acetate (AmAc) concentrations following a 10-minute incubation at 20 °C.**

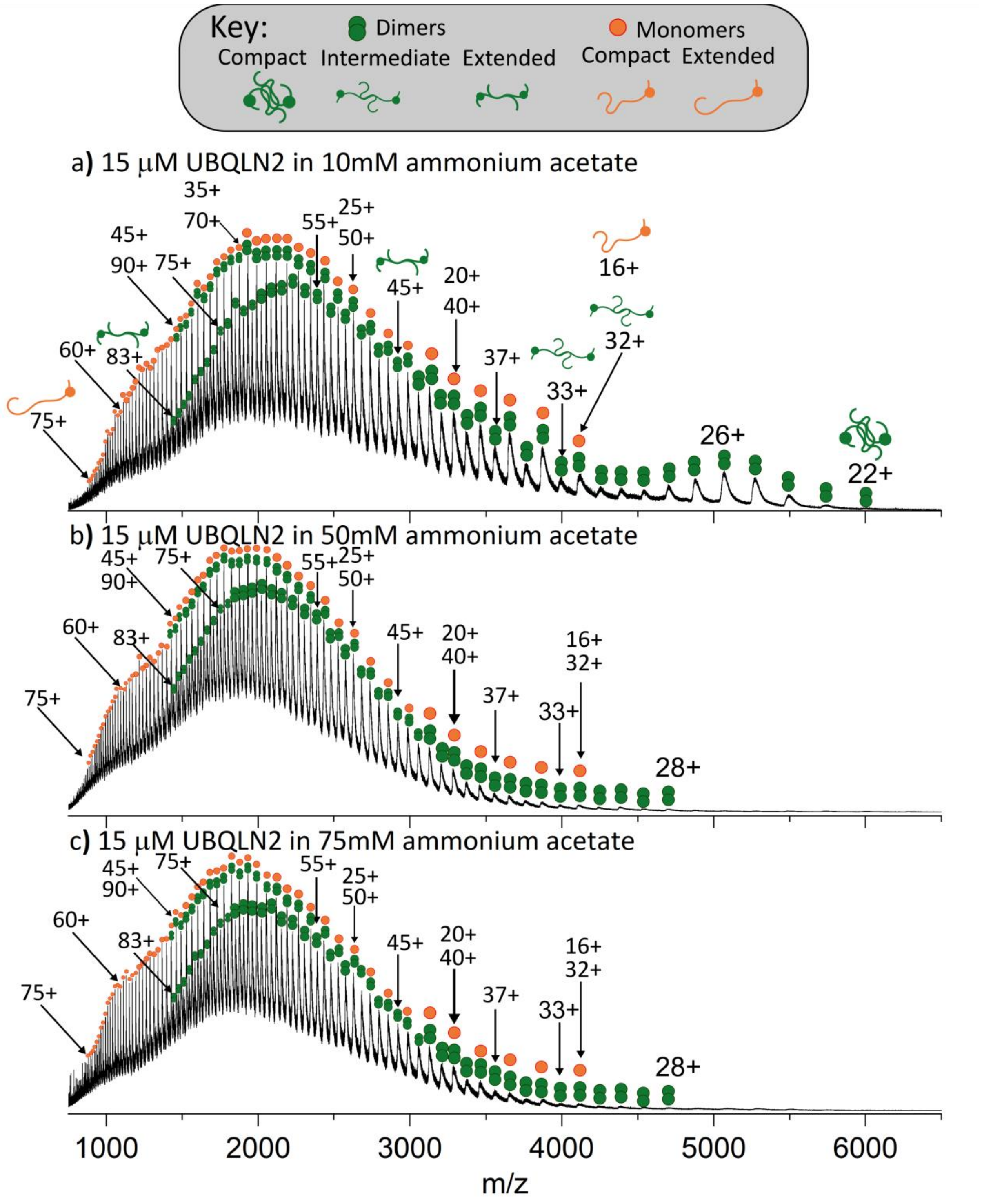


**Figure S2: Fully annotated native mass spectra of FL-UBQLN2 at a) 10mM, b) 50mM and c) 75mM AmAc.**

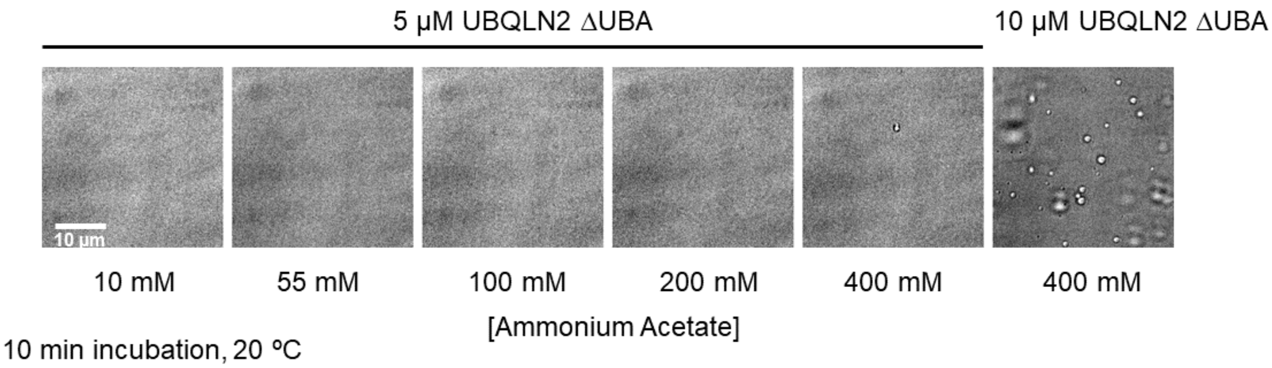


**Figure S3: Brightfield microscopy of UBQLN2-ΔUBA showing it is fully soluble below 400mM AmAc at 5μM but undergoes LLPS at higher protein concentrations following a 10-minute incubation at 20 °C.**

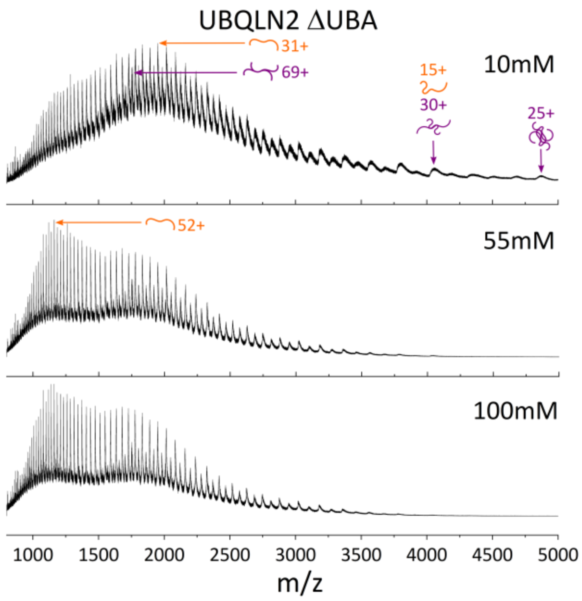


**Figure S4: Native mass spectra of UBQLN2-ΔUBA (5μM), sprayed from 10mM (top), 55mM (middle) and 100mM (bottom) AmAc concentration. The loss of compact dimers follows the same trend as FL-UBQLN2.**

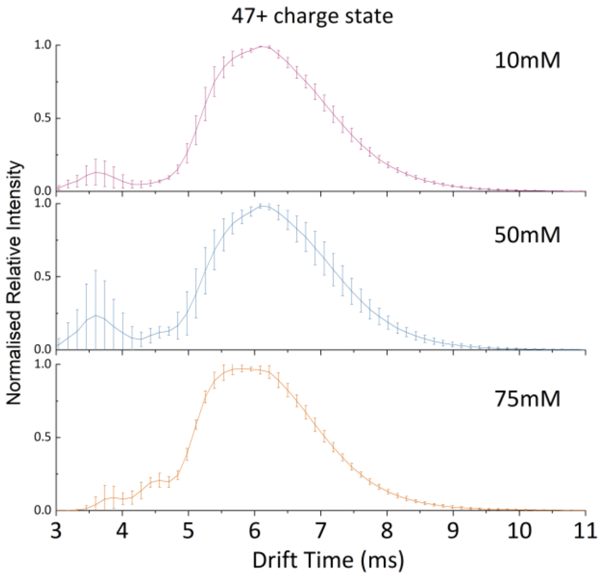


**Figure S5: Ion mobility spectra at 10mM (top), 50mM (middle) and 75mM (bottom) for 47+ dimer charge state of FL-UBQLN2. Changes in the AmAc concentration from which the protein is sprayed does not majorly alter the arrival time distribution of the more extended conformations.**

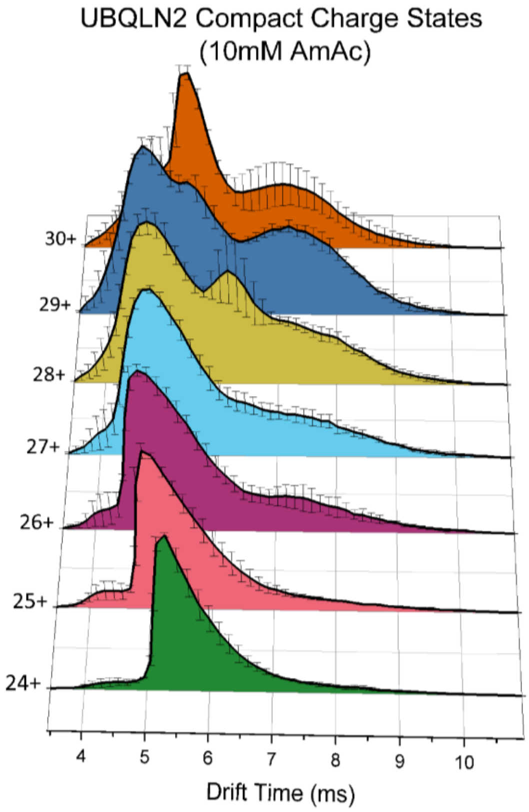


**Figure S6: Arrival time distributions of charges states 24+ to 30+, corresponding to compact charge states, at 10mM AmAc. FL-UBQLN2 exists as a compact conformation, centred around 5.5 ms in the 24+ charge state which arrives earlier as charge increases. A new, more extended conformation can be seen from the 26+ charge state onwards at around 7.5 ms which increases in relative intensity as charge increases. The peaks at 6.5 ms in the 28+ and 5.5ms 30+ charge states correspond to monomeric FL-UBQLN2.**

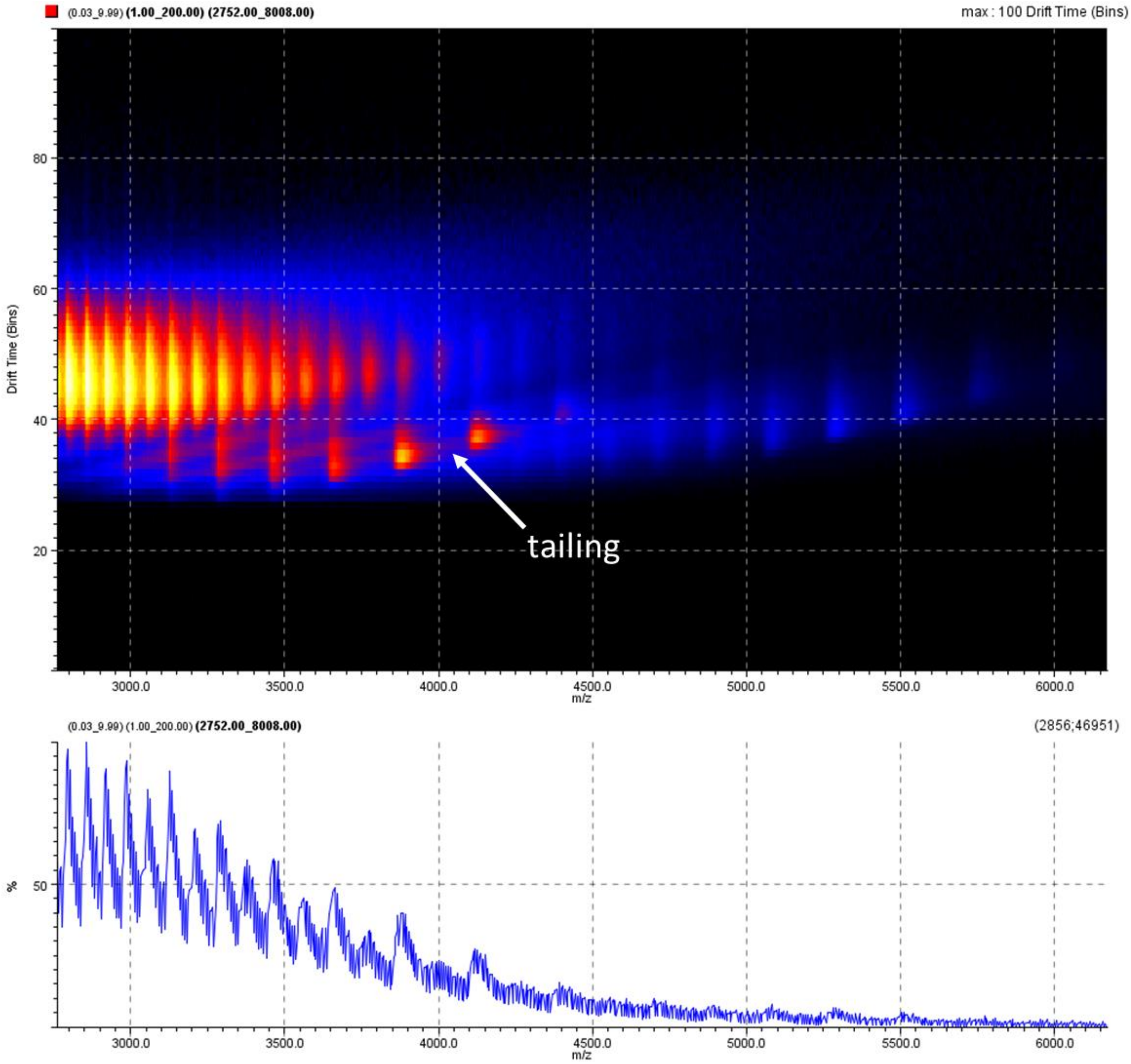


**Figure S7: IMS data of FL-UBQLN2 shown in Driftscope heatmap format with square root intensity averaging for one representative file collected at 10mM AmAc with manual quad profile of m/z 3500 (see method for more details) to confirm absence of higher order species in the space beneath charge states in the mass spectrum.**

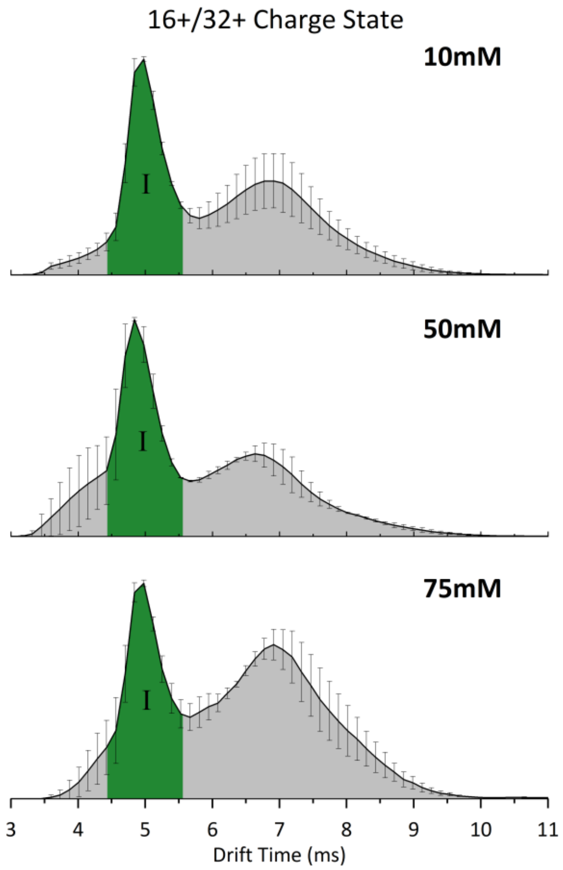


**Figure S8: IM spectra of the FL-UBQLN2 16+ monomer, 32+ dimer sprayed from AmAc concentrations of 10 mM (top), 50 mM (middle) and 75 mM (bottom). Sharp profile for peak I, shown in green, can be attributed to the monomer as it is not present in the 33+ dimer charge state. This peak is not altered by an increase in AmAc concentration.**

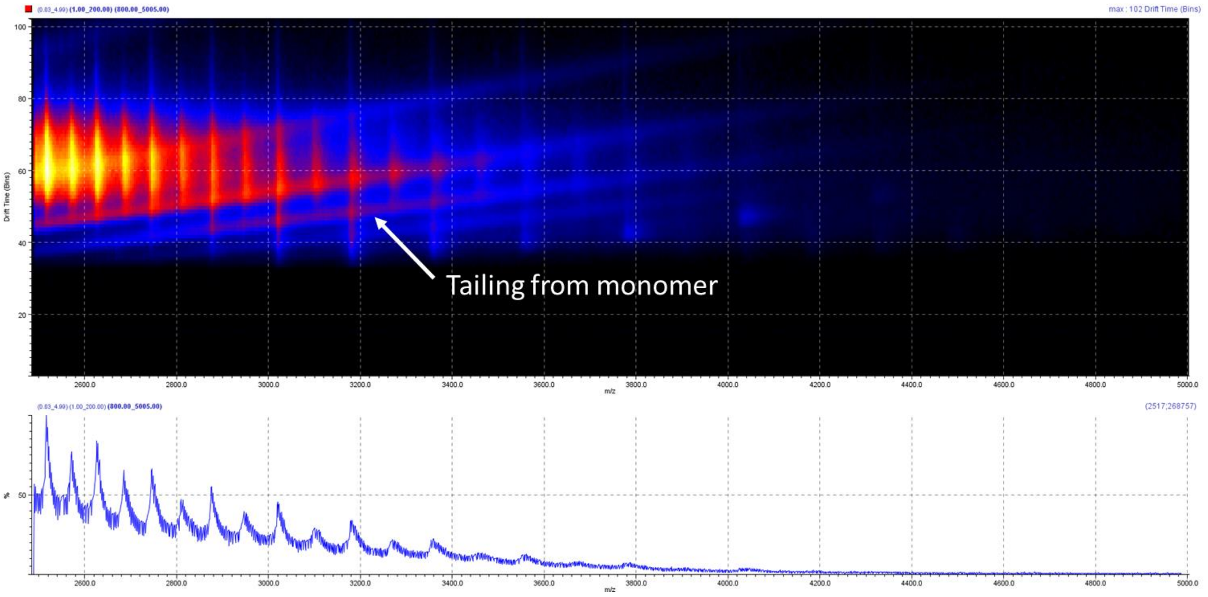


**Figure S9: IMS data of UBQLN2-ΔUBA shown in Driftscope heatmap format with square root intensity averaging for one representative file collected at 55mM AmAc to show 'tailing' of the monomers into odd-charged dimers in *m/z* space.**

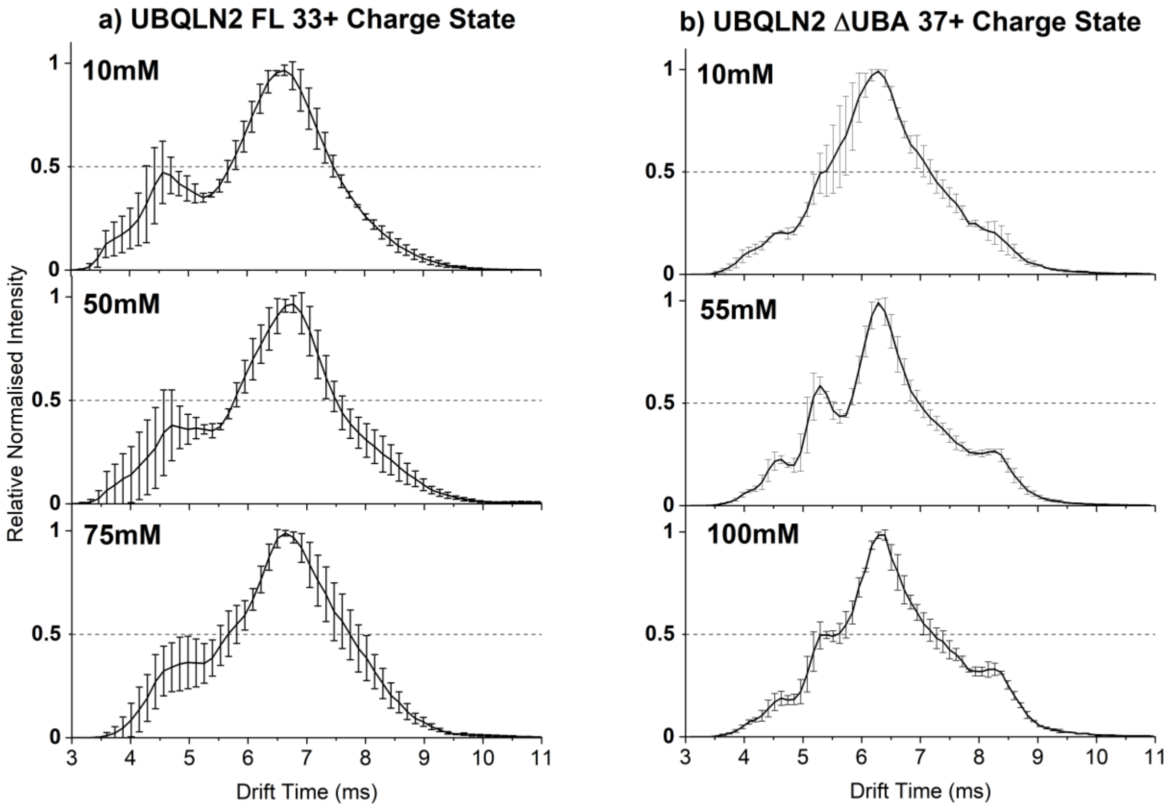


**Figure S10: IM spectra of a) UBQLN2 FL 33+ Charge State and b) UBQLN2-ΔUBA 37+ charge state without fitted peaks present.**

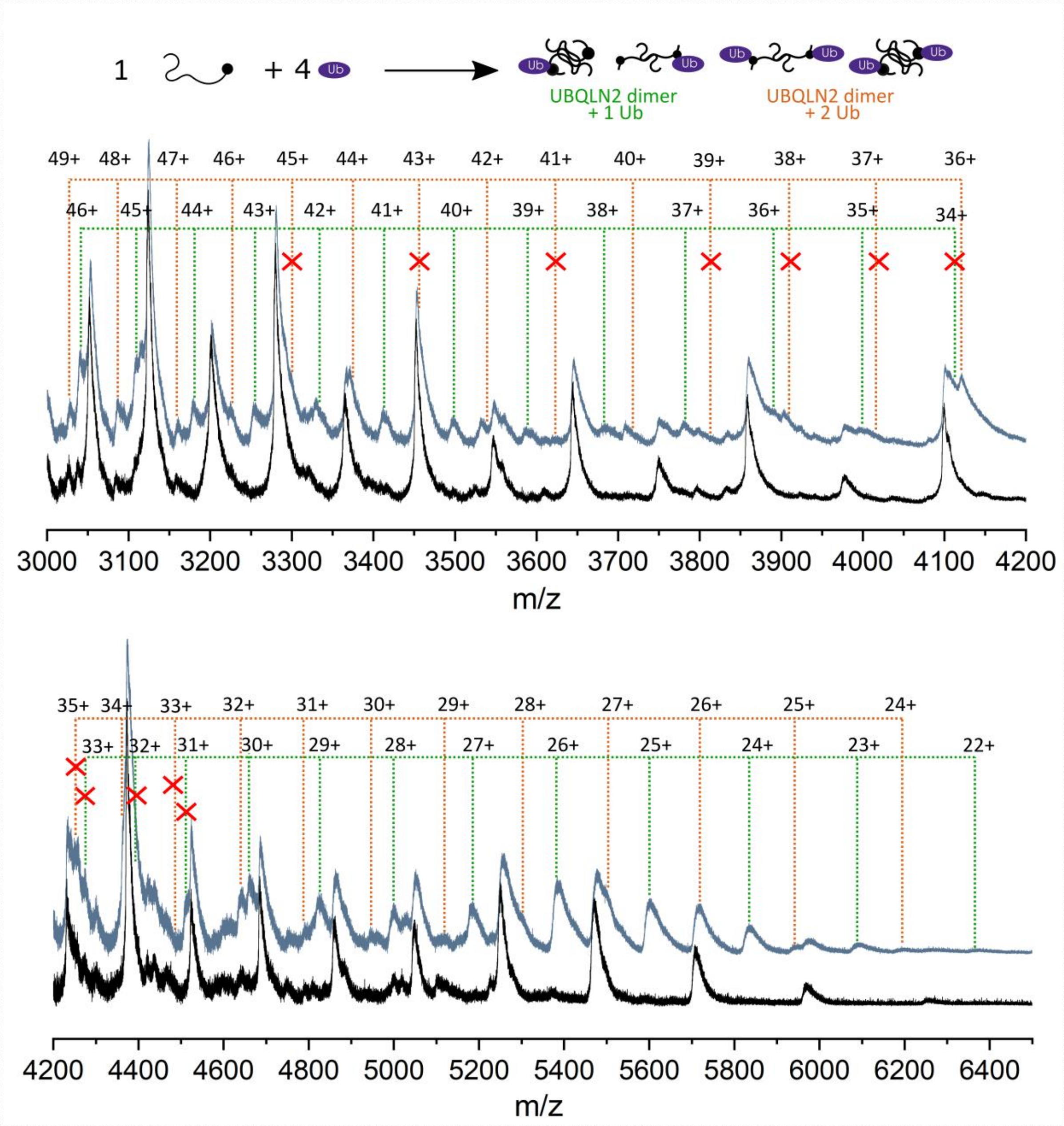


**Figure S11: Native mass spectrum of FL-UBQLN2 alone (black) and resulting from 1:4 molar ratio of UBQLN2:Ub (blue) overlayed and offset vertically to allow comparison of charge states in the region 3000-4200 m/z (top) and 4200-6500 m/z (bottom). All regions were baseline subtracted using Subtract Baseline function on OriginPro (see methods for more details). Labels marked with red crosses indicate charge states which are not well resolved in the mass spectrum and so m/z has been estimated, and calibrated CCS values for these charge states are not shown in figure 5.**

**Table S1 UBQLN2 Monomer Charge States**

| Z | m/z | Z | m/z | Z | m/z | Z | m/z | Z | m/z |
|---|---|---|---|---|---|---|---|---|---|
| ***16*** | 4104.4 | ***28*** | 2345.8 | ***40*** | 1642.4 | ***52*** | 1263.6 | ***64*** | 1026.9 |
| ***17*** | 3863.0 | ***29*** | 2264.9 | ***41*** | 1602.3 | ***53*** | 1239.8 | ***65*** | 1011.1 |
| ***18*** | 3648.5 | ***30*** | 2189.5 | ***42*** | 1564.2 | ***54*** | 1216.8 | ***66*** | 996.4 |
| ***19*** | 3456.5 | ***31*** | 2118.9 | ***43*** | 1527.8 | ***55*** | 1194.7 | ***67*** | 981.5 |
| ***20*** | 3283.7 | ***32*** | 2052.7 | ***44*** | 1493.1 | ***56*** | 1173.4 | ***68*** | 967.1 |
| ***21*** | 3127.4 | ***33*** | 1990.5 | ***45*** | 1460.0 | ***57*** | 1152.8 | ***69*** | 953.1 |
| ***22*** | 2985.3 | ***34*** | 1932.0 | ***46*** | 1428.3 | ***58*** | 1133.0 | ***70*** | 939.5 |
| ***23*** | 2855.5 | ***35*** | 1876.8 | ***47*** | 1397.9 | ***59*** | 1113.8 | ***71*** | 926.3 |
| ***24*** | 2736.6 | ***36*** | 1824.7 | ***48*** | 1368.8 | ***60*** | 1095.2 | ***72*** | 913.5 |
| ***25*** | 2627.2 | ***37*** | 1775.4 | ***49*** | 1340.9 | ***61*** | 1077.3 | ***73*** | 901.0 |
| ***26*** | 2526.2 | ***38*** | 1728.7 | ***50*** | 1314.1 | ***62*** | 1059.9 | ***74*** | 888.8 |
| ***27*** | 2432.6 | ***39*** | 1684.4 | ***51*** | 1288.3 | ***63*** | 1043.1 | ***75*** | 877.0 |

**Table S2 UBQLN2 Dimer Charge States**

| Z | m/z | Z | m/z | Z | m/z | Z | m/z | Z | m/z |
|---|---|---|---|---|---|---|---|---|---|
| ***22*** | 5969.6 | ***35*** | 3752.7 | ***48*** | 2736.6 | ***61*** | 2153.6 | ***74*** | 1775.4 |
| ***23*** | 5710.1 | ***36*** | 3648.5 | ***49*** | 2680.8 | ***62*** | 2118.9 | ***75*** | 1751.8 |
| ***24*** | 5472.2 | ***37*** | 3549.9 | ***50*** | 2627.2 | ***63*** | 2085.3 | ***76*** | 1728.7 |
| ***25*** | 5253.3 | ***38*** | 3456.5 | ***51*** | 2575.7 | ***64*** | 2052.7 | ***77*** | 1706.3 |
| ***26*** | 5051.3 | ***39*** | 3367.9 | ***52*** | 2526.2 | ***65*** | 2021.1 | ***78*** | 1684.4 |
| ***27*** | 4864.3 | ***40*** | 3283.7 | ***53*** | 2478.5 | ***66*** | 1990.5 | ***79*** | 1663.1 |
| ***28*** | 4690.6 | ***41*** | 3203.6 | ***54*** | 2432.6 | ***67*** | 1960.8 | ***80*** | 1642.4 |
| ***29*** | 4528.9 | ***42*** | 3127.4 | ***55*** | 2388.4 | ***68*** | 1932.0 | ***81*** | 1622.1 |
| ***30*** | 4377.9 | ***43*** | 3054.7 | ***56*** | 2345.8 | ***69*** | 1904.0 | ***82*** | 1602.3 |
| ***31*** | 4236.8 | ***44*** | 2985.3 | ***57*** | 2304.7 | ***70*** | 1876.8 | ***83*** | 1583.0 |
| ***32*** | 4104.4 | ***45*** | 2919.0 | ***58*** | 2264.9 | ***71*** | 1850.4 | | |
| ***33*** | 3980.0 | ***46*** | 2855.5 | ***59*** | 2226.6 | ***72*** | 1824.7 | | |
| ***34*** | 3863.0 | ***47*** | 2794.8 | ***60*** | 2189.5 | ***73*** | 1799.7 | | |

**Table S3 UBQLN2:Ub 2:1 complex Charge States**

| Z | m/z | Z | m/z | Z | m/z |
|---|---|---|---|---|---|
| ***22*** | 6362.7 | ***34*** | 4117.4 | ***46*** | 3043.5 |
| ***23*** | 6086.1 | ***35*** | 3999.8 | | |
| ***24*** | 5832.5 | ***36*** | 3888.7 | | |
| ***25*** | 5599.3 | ***37*** | 3783.6 | | |
| ***26*** | 5384.0 | ***38*** | 3684.1 | | |
| ***27*** | 5184.6 | ***39*** | 3589.6 | | |
| ***28*** | 4999.5 | ***40*** | 3499.9 | | |
| ***29*** | 4827.1 | ***41*** | 3414.6 | | |
| ***30*** | 4666.2 | ***42*** | 3333.3 | | |
| ***31*** | 4515.7 | ***43*** | 3255.8 | | |
| ***32*** | 4374.7 | ***44*** | 3181.8 | | |
| ***33*** | 4242.1 | ***45*** | 3111.2 | | |

**Table S4 UBQLN2:Ub 2:2 complex Charge States**

| | | | | | |
|---|---|---|---|---|---|
| ***24*** | 6186.0 | ***34*** | 4366.9 | ***44*** | 3374.6 |
| ***25*** | 5938.6 | ***35*** | 4242.2 | ***45*** | 3299.7 |
| ***26*** | 5710.2 | ***36*** | 4124.3 | ***46*** | 3228.0 |
| ***27*** | 5498.8 | ***37*** | 4012.9 | ***47*** | 3159.3 |
| ***28*** | 5302.4 | ***38*** | 3907.3 | ***48*** | 3093.5 |
| ***29*** | 5119.6 | ***39*** | 3807.2 | ***49*** | 3030.4 |
| ***30*** | 4949.0 | ***40*** | 3712.0 | | |
| ***31*** | 4789.4 | ***41*** | 3621.5 | | |
| ***32*** | 4639.8 | ***42*** | 3535.3 | | |
| ***33*** | 4499.2 | ***43*** | 3453.1 | | |

Red = unresolved in mass spectrum, as labelled with X in Figure S7.
